# CRISPR/Cas9 mutants delineate roles of *Populus FT* and *TFL1*/*CEN*/*BFT* family members in growth, dormancy release and flowering

**DOI:** 10.1101/2022.08.10.503493

**Authors:** Xiaoyan Sheng, R. Ayeshan Mahendra, Chieh-Ting Wang, Amy M. Brunner

**Affiliations:** Department of Forest Resources and Environmental Conservation, Virginia Tech, Blacksburg, VA 24061, USA

**Author notes:** Corresponding author A.M. Brunner. These authors contributed equally (Xiaoyan Sheng, R. Ayeshan Mahendra).

**Keywords:** Poplar, floral transition, meristem identity, seasonal signals, PHOSPHATIDYLETHANOLAMINE-BINDING PROTEIN

## Abstract

Vegetative and reproductive phase change and phenology are economically and ecologically important traits. Trees typically require several years of growth before flowering and once mature, seasonal control of the transition to flowering and flower development is necessary to maintain vegetative meristems and for reproductive success. Members of two related gene subfamilies, *FLOWERING LOCUST (FT)* and *TERMINAL FLOWER1 (TFL1)/CENTRORADIALIS (CEN)/BROTHER OF FT AND TFL1 (BFT)*, have antagonistic roles in flowering in diverse species and roles in vegetative phenology in trees, but many details of their functions in trees have yet to be resolved. Here, we used CRISPR/Cas9 to generate single and double mutants involving the five *Populus FT* and *TFL1/CEN/BFT* genes. *ft1* mutants exhibited wild-type-like phenotypes in long days and short days, but after chilling to release dormancy showed delayed bud flush and GA_3_ could compensate for the *ft1* mutation. After rooting and generating some phytomers in tissue culture, both *cen1* and *cen1ft1* mutants produced terminal as well as axillary flowers, indicating that the *cen1* flowering phenotype is independent of *FT1*. Some axillary meristems initially generated phytomers and in potted plants, the timing of flowering in these shoots correlated with upregulation of *FT2* in maturing leaves, suggesting that, in long days, *CEN1* antagonizes *FT2* promotion of flowering but enables *FT2* promotion of shoot growth by maintaining indeterminacy of the shoot apical meristem. *CEN1* showed distinct circannual expression patterns in vegetative and reproductive tissues and comparison with the expression patterns of *FT1* and *FT2* suggest that the relative levels of *CEN1* compared to *FT1* and *FT2* regulate multiple phases of vegetative and reproductive seasonal development.

## Introduction

The floral transition in diverse plants and vegetative phenology of temperate and boreal trees are regulated by similar environmental signals and recognition of this similarity led to the discovery that *Populus FLOWERING LOCUS T2* (*FT2*) regulates short day (SD)-induced growth cessation and bud set (Böhlenius et al. 2006; Hsu et al. 2011). Recently, mutagenesis of *FT2* showed its essential role in maintaining shoot growth in long days (LDs; Andre et al. 2022a; Gomez-Soto et al. 2021). FT2 is a member of the PHOSPHATIDYLETHANOLAMINE-BINDING PROTEIN (PEBP) family that in angiosperms consists of three major clades, here named for family members: FT, **T**ERMINAL FLOWER1 (TFL1)/**C**ENTRORADIALIS(CEN)/ **B**ROTHER OF FT AND TFL1 (BFT) (TCB clade) and MOTHER OF FT AND TFL1 (MFT) (Bennett and Dixon 2021). The balance between florigen FT and indeterminacy-promoting TCB members is a broadly conserved mechanism to control flowering time and plant architecture; however, gene duplication and evolution have resulted in functional diversification among paralogs and orthologs (reviewed in Jin et al. 2021; Wickland and Hanzawa 2015). Both selection of natural alleles of PEBP genes and more recently, targeted mutations, have resulted in desirable flowering and architecture phenotypes in crops (Eshed and Lippman 2019); and in *Populus*, genetic variation in *FT1* and *FT2* has been associated with bud phenology (Evans et al. 2014; Wang et al. 2018).

Vegetative phase change, typically changes in leaf characteristics, and reproductive phase change — the acquisition of competency to respond to flower-promoting signals — are coordinated in Arabidopsis, but these occur at very different times in trees such as *Eucalyptus* and *Acacia* (reviewed in Brunner et al. 2017). In *Populus*, vegetative phase change occurs within a few months of growth and, as in several herbaceous species, this is regulated by miR156 (Lawrence et al. 2021). The genetic mechanisms controlling reproductive phase change in trees remain elusive and in adult trees, only a subset of meristems commit to flowering during a limited seasonal time, suggesting that there is a recurring seasonal acquisition of competency to flower in adult trees. In *Arabis alpina*, a perennial polycarpic relative of Arabidopsis, vernalization is required for flowering but if plants are too young when vernalized, flowering does not occur. A *TFL1* ortholog was shown to have role in this age-dependent response to vernalization as well as in polycarpy (Wang et al. 2011).

The circannual expression patterns of *Populus FT1* and *FT2* are divergent with *FT1* predominately expressed in winter and *FT2* during the growing season (Hsu et al. 2011). Their encoded proteins show more subtle differences, particularly in regards to flowering promotion. Under the direction of a heat-inducible promoter, *FT1* but not *FT2* induces flowering and when driven by the 35S promoter, *FT1* induces wild-type (WT)-like catkins, while *FT2* induces individual flowers (Böhlenius et al. 2006; Hsu et al. 2011; Hsu et al. 2006). These characteristics of *FT1* combined with detailed study of shoot morphogenesis showing that inflorescences developed in the axils of late preformed leaves (LPL; (Yuceer et al. 2003) indicated that *FT1* promotes the floral transition (Hsu et al. 2011). Controlled environment studies showed that following SD-induced dormancy, *FT1* and GA biosynthesis and signaling genes are upregulated during the chilling treatment to release dormancy (Rinne et al. 2011). Moreover, lipid body (LB)-associated glucan hydrolase family 17 (GH17) enzymes thought to have a role in opening of plasmodesmata were also upregulated during chilling as well as by GA_3_, suggesting that *FT1* might promote dormancy release by promoting GA_3_ synthesis and signaling.

*Populus* members of the TCB clade include two co-orthologs of *CEN* and a *BFT* ortholog (Figure 1a). RNAi-mediated downregulation of *CEN1/CEN2* induced an earlier first onset of flowering at the normal seasonal time and axillary position, but it took three years of growth before the RNAi trees were consistently flowering and producing many catkins (Mohamed et al. 2010). Additionally, bud flush was delayed in *35S:CEN1* trees in field environments and controlled environment studies of RNAi and overexpression transgenics showed opposite effects on dormancy release. Whereas the aforementioned studies indicate that *Populus FT* and *TCB* clade members have roles in both flowering and vegetative phenology, much remains to be delineated. For loss-of function, past studies had to rely on RNAi-mediated silencing that does not discriminate among paralogs (e.g., Klocko et al. 2016), limiting understanding of endogenous functions. Here, we report the phenotypes induced in single and double CRISPR/Cas9-mediated *Populus* mutants of *FT* and *TCB* clade members. Our findings include that among the *TCB* members, *CEN1* is essential to prevent flowering under growth-conducive conditions and that *FT1* promotes dormancy release/bud flush. In addition, circannual expression of *CEN1* and *CEN2* in multiple tissues that could be directly correlated with previously reported circannual expression of *FT1, FT2* as well as *FLOWERING LOCUS D-LIKE2* (*FDL2*) (Hsu et al. 2011; Sheng et al. 2022) further suggest that the relative levels of *CEN1* and *FT1* or *FT2* regulate multiple stages of vegetative and flowering phenology by acting in different tissues at different seasonal times.

**Figure 1.**
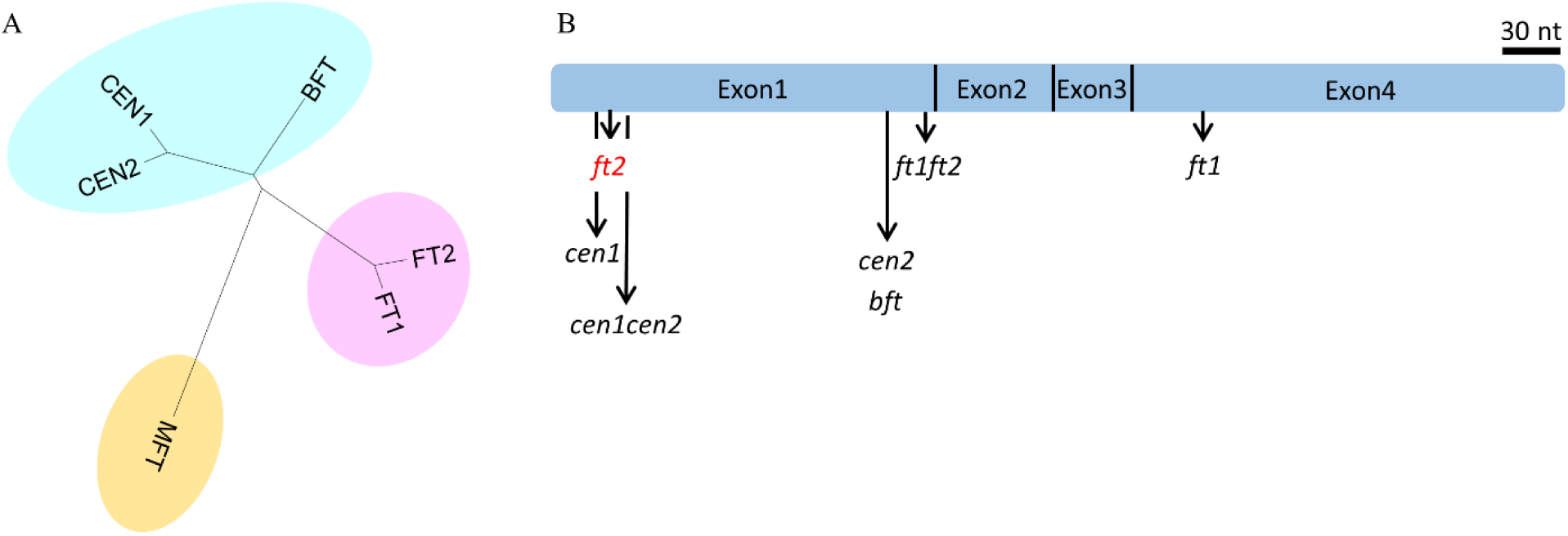
*Populus* PEBP family mutants. (A) Neighbor-Joining tree of *Populus* PEBP family proteins with the three major angiosperm clades in different colors. (B) Schematic of the coding sequence of *FT/CEN/BFT* showing locations of gRNA target sites and resulting mutants (all that were confirmed had 1-2 bp indels). Red type denotes that no shoots regenerated following transformation with the construct targeting *FT2*. Not shown are *cen1ft1* mutants that were generated by introducing the construct targeting *CEN1* into the homozygous *ft1-3* mutant. Additional details of the mutants, including target site sequences are provided in Figure S2.

## Materials and Methods

### CRISPR/Cas9-mediated mutants

Guide RNAs (gRNAs) were designed according to the steps outlined in (Li et al. 2013) and the selected 23-base pair (bp) sequences ending with the protospacer-adjacent motif (PAM) were checked for target specificity by BLAST against the custom VariantDB for the transformation clone *Populus tremula* × *alba* INRA 717-1B4 (Xue et al., 2015). Each of the target site sequences was introduced into *Arabidopsis* U6 promoter-gRNA cassette cloned into pGEM-T Easy plasmid (Promega) using two-step overlapping PCR (Figure S1); primers used to generate the gRNAs are listed in Table S1. The gRNAs were excised and inserted at the Pme1 site of the pMOA33-UBQ Cas9 OCS binary vector (Peterson et al. 2016); Figure S1B). All constructs were introduced into *Agrobacterium tumefaciens* strain GV3101 and transformed into *P. tremula* x *P. alba* clone INRA 717-1B4, hereafter referred to as wild-type (WT), as previously described (Meilan and Ma 2006). To generate *cen1ft1* mutants, the gRNA targeting *CEN1* was introduced into the homozygous *ft1-3* mutant. To determine genetic mutation patterns of the target gene in transgenic trees, the genomic region spanning the gRNA target site was PCR-amplified (primers are listed in Table S1) from WT and gene-edited tree genomic DNA using iProof GC master mix (Bio-Rad) and sequenced. Mutations in select events were confirmed by cloning PCR fragments into pGEM-T Easy and sequencing from 10 colonies. For the closest paralogs (i.e., *FT1/FT2* and *CEN1/CEN2*), we also amplified the corresponding region in the non-target paralog to confirm the absence of a mutation.

### Environmental treatments

WT and *ft1* mutant plants were propagated *in vitro*. Rooted plantlets were transferred from tissue culture to soil (Promix B, Canada) and acclimated in a LD^16hr^ growth chamber at 22°C day/20 °C night. After acclimation, plants were transferred to two-gallon pots and 48 g of Osmocote Plus 15-9-12 fertilizer/pot added approximately 3 weeks after transfer. Plants were grown under LD^16hr^ at 22°C day/20 °C night until reaching heights of 65 to 70 cm. Growth chamber settings were then changed to SD^8hr^ with the same temperatures (22°C day/20 °C night) to induce growth cessation and dormancy. After 6 weeks inSD^8h^, we lowered temperatures to 15°C day /12 °C night for 2 weeks, a transitional phase for acclimation to chilling temperatures, followed by 10°C day /8 °C night temperatures for 6 weeks to release dormancy. Finally, the plants were transferred to a LD^16hr^ and warm temperature (22/20 °C) greenhouse to assess dormancy release by scoring time of bud flush. To study the growth phenotypes of WT and *bft* mutant trees to nutrient depletion, no fertilizer was added after transfer to two-gallon pots containing Promix B (Canada), and growth and bud stage were monitored in a LD^16hr^ greenhouse.

### Bud-internode unit assay for bud flush

A bud-internode unit assay for bud flush was modified from the method previously described (Rinne et al. 2011). Before the experiment, mutant and WT plants were exposed to 8 weeks of SD^8h^ followed by 6 weeks of chilling temperatures as described in the preceding paragraph. The stem of each plant was then cut into segments including an axillary bud and 1- to 2-cm-long internodal segment both above and below the bud. The apical bud-internode unit along with each of the next 23 axillary bud-internode units was placed in a well with 1 ml of water as treatment control or 1 µM GA_3_ (Sigma-Aldrich) solution. To uncover any effects of bud position on the shoot, bud-internode units were placed in order from left to right, in each row of a 24-well cell culture plate. The plates were incubated in forcing conditions of continuous light at room temperature to monitor the time of visible bud flush.

### Gene expression analysis

Leaf samples were collected from the soil grown WT plants and in vitro grown WT plants to analyze the *FT1* and *FT2* gene expression. Leaf Plastrochron Index (LPI) (Larson and Isebrands 1971) where the first leaf with a lamina longer than 1 cm was considered as LPI 1, was used to standardize leaf developmental stages across plants. Total RNA was extracted from the collected leaf samples using the RNeasy Plant Mini Kit (Qiagen) and was treated with RNase*-*Free DNase Set (Qiagen) as described by (Brunner et al. 2004). Each cDNA was synthesized from 3.0 μg total RNA and an oligo (dT) primer using the High Capacity cDNA reverse transcription kit (Applied Biosystems), according to the manufacturer’s protocol. cDNA samples were diluted ten times with nuclease free water prior to the quantitative real time PCR. Quantitative real time PCR reactions were done in ABI PRISM™ 7500 Real-Time PCR system using Power SYBR Green PCR Master Mix kit (Applied Biosystems), with three replicates per RNA sample. The PCR program was set up to perform an initial incubation at 95°C for 10 min, followed by 95°C for 15 s, and 60°C for 1 min, for a total of 40 cycles. We used an ubiquitin gene (*UBQ2*) as an internal reference (Mohamed et al. 2010), and normalized the Ct values across plates, determining relative quantities using comparative Ct method (2^−Δ·ΔCt^) as previously described (Livak and Schmittgen 2001).

For study of *CEN1* and *CEN2* circannual expression, we used the same *P. deltoides* samples and parameters for qRT-PCR that were previously used to study *FT1* and *FT2* expression (Hsu et al. 2011) and samples are summarized in Supplementary Table S2. We used *TIP41L* (Gutierrez et al. 2008) as an internal reference and calculated relative quantities as described in the preceding paragraph. All primer sequences and gene IDs are listed in Table S1.

## Results

### ft1 mutants appear wild-type under LDs whereas ft1ft2 mutants set terminal buds

To differentiate the functions of *FT1* and *FT2*, we designed gRNA specifically targeting each of them as well as a gRNA that targeted both paralogs (Figure 1, Figure S2). We were unable to regenerate any plants following transformation with the *FT2*-specific gRNA, but we did generate *ft1ft2* mutants. This could be due to mutation at the *FT2-*specfic site having a stronger effect on regeneration than a mutation at the shared *FT1/FT2* target site. However, most transformants with the gRNA targeting both *FT1* and *FT2* failed to regenerate shoots. Only a few short shoots were regenerated, and some of these set buds while still on shoot elongation media plates under LD growing conditions. We were able to root three independent events that harbored 1-2 bp indels (Figure S2). The mutants had extremely short internodes and set terminal bud when grown in magenta boxes for 6 weeks under LD^16hr^ (Figure 2a).

**Figure 2.**
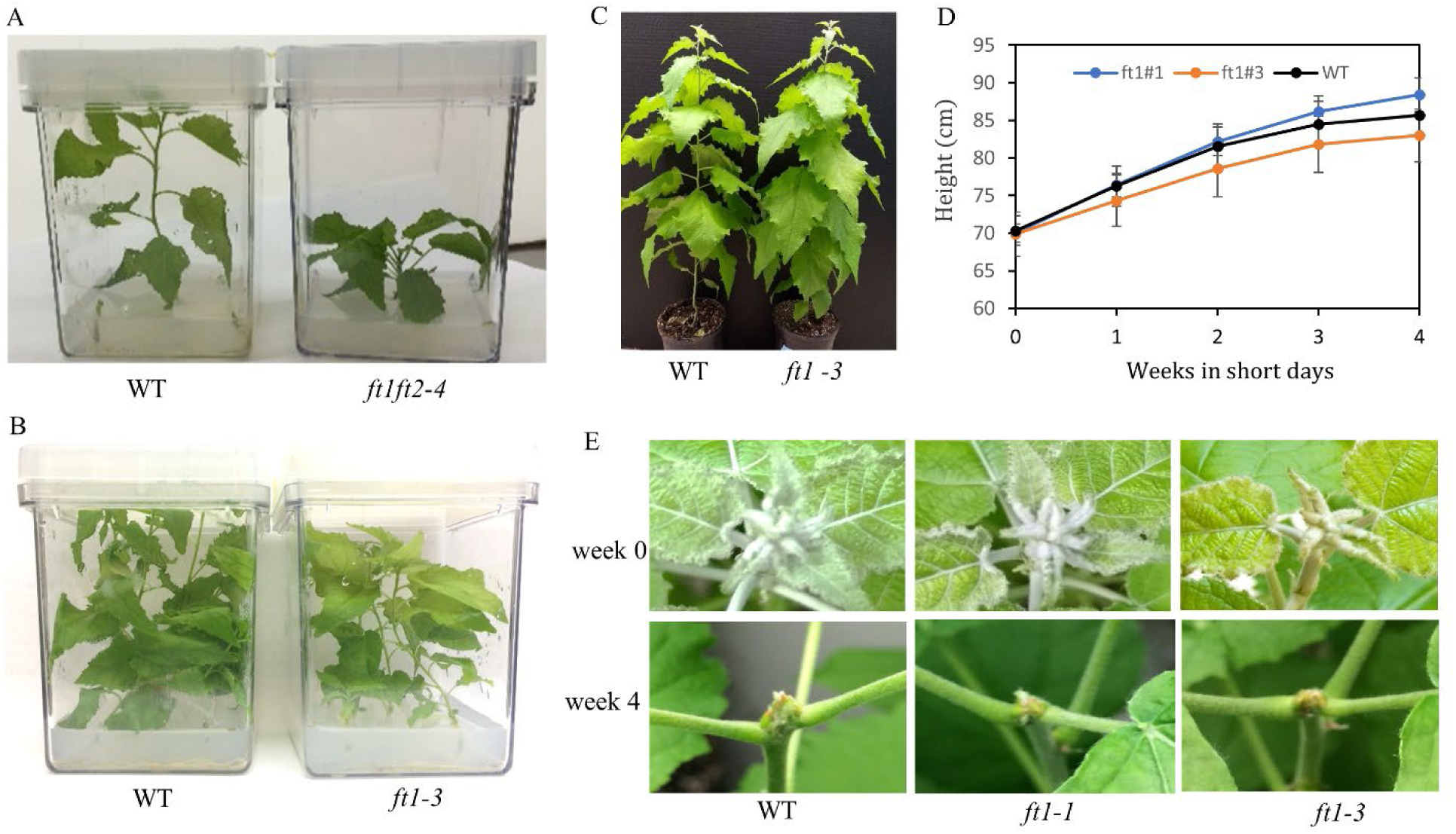
*ft1ft2* mutants set terminal buds in long days whereas *ft1* mutants show WT-like growth in both long days and short days. (A) Six-week-old *in vitro* plants grown in LDs. The *ft1ft 2-4* plant has set a terminal bud. Representative WT and *ft1-3* plants after two months growth in LDs in tissue culture (B) and soil (C). (D) Progression of SD-induced growth cessation in WT and *ft1-3* mutants. Means ± SE (n = 5). (E) Birds eye view of shoot apex of representative WT and *ft1* plants after 3 months in LDs (week 0) followed by 4 weeks of SDs.

We regenerated multiple events and studied two independent events harboring heterozygous or homozygous frame shift indels in the *FT1* locus (Figure 1, Figure S2). No phenotypic alterations were noted in either *ft1* mutant compared to WT when grown under LD^16hr^ in tissue culture or soil (Figure 2 B, C).

### ft1 mutants show delayed bud flush phenology

Both *ft1*-1 and *ft1*-3 mutants ceased growth at the same time as WT after they were transferred to SD^8hr^ conditions (Figure 2d). The mutants also showed no differences in progression of terminal bud formation from WT, and all genotypes completed bud set after exposure to 4 weeks of SD^8hr^ (Figure 2E). The trees continued to be exposed to SD^8hr^ for a total of 8 weeks to induce dormancy, and then temperatures were lowered to promote dormancy release (Figure 3A). After 6 weeks in chilling temperatures, trees were transferred to a LD^16hr^ and warm temperature greenhouse. Terminal and axillary buds on all WT plants started to swell after 7-10 days in the greenhouse, and internode elongation began after preformed leaves emerged and unfolded within 11-15 days. In contrast, all *ft1*-1 and *ft1*-3 plants remained quiescent (Figure 3B). However, terminal buds and a few axillary buds on the four *ft1*-*1* mutant plants flushed after 4 weeks in the greenhouse, while buds of *ft1*-3 plants flushed after 6 weeks in the greenhouse.

**Figure 3.**
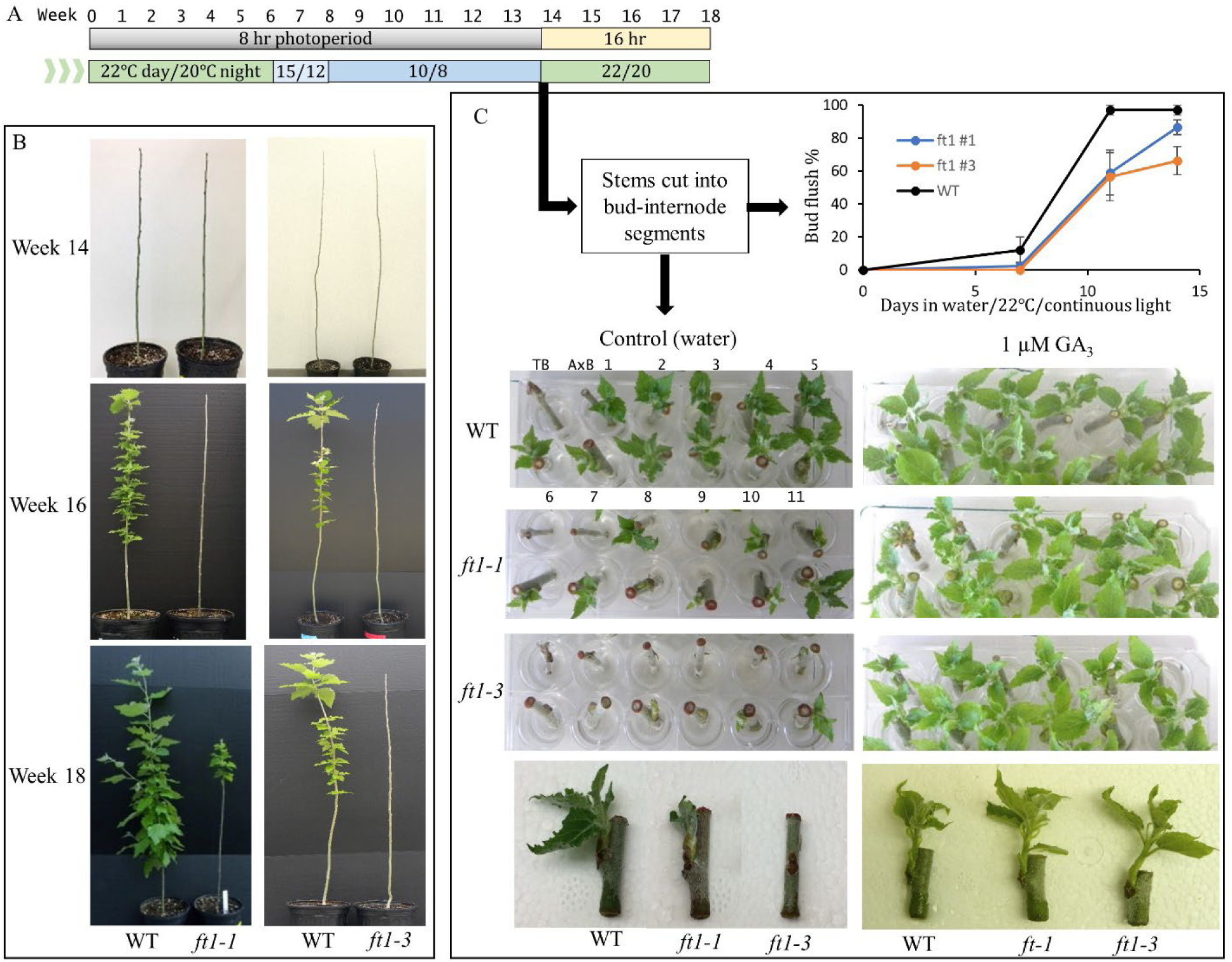
Bud flush is delayed in *ft1* mutants. (A) Daylength and temperature regime to induce growth cessation and dormancy (SD, warm temperature), dormancy release (chilling temperature) and bud flush (LD, warm temperature). (B) Representative WT and *ft1* trees at post-chilling timepoints. (C) Detached bud-internode assay showing that GA_3_ compensates for *ft1* mutation. After the chilling phase, stems were cut and segments were incubated at room temperature with continuous light. Bud-internode segments are ordered in all plates as indicated for the WT control. TB, terminal bud with internode; AxB (axillary bud-internode segments) from 1 (first bud below the TB) to 11. Bottom photographs show axillary bud-internode segment 10.

To further understand the role of *FT1* in dormancy release, we adopted a culture system of bud-internode units (Rinne et al. 2011) to assess the impact of chilling temperatures and GA-feeding. WT, *ft1*-*1* and *ft1*-*3* trees were exposed to 8 weeks of SD^8hr^ followed by 6 weeks of chilling temperatures (Figure 3A). The stems of each plant were cut into terminal bud-internode and axillary bud-internode segments, and each segment was placed in a well with either water (control treatment) or a GA_3_ (1 µM) solution at room temperature and continuous light. In the control treatment, except for the terminal bud, the top 10 axillary buds of WT flushed within 7-14 days (Figure 3C). Both *ft1*-1 and *ft1*-3 mutants showed delayed bud flush when incubated in water (Figure 3C). However, all the axillary buds from the lower region of the stem, (buds from positions 11 to 23) of the *ft1* mutants and WT flushed after 7 days in water (Figure S3). This suggests that the impact of *FT1* on dormancy release of axillary buds might reduce with increasing distance from the apical bud.

In many plants, application of GA can substitute for a prolonged exposure to chilling temperatures (Saure 1985). In *P. tremula* × *P. tremuloides*, chilling induced the upregulation of GA biosynthesis genes and promoted the reopening of plasmodesmata and the response to growth promoting signals (Rinne et al. 2011). All tested apical and the axillary buds from both *ft1*-1 and *ft1*-3 flushed at the same time as WT plants, within 5-7 days, when incubated in the GA_3_ solution (Figure 4), consistent with *FT1* promoting dormancy release by activating GA synthesis, transport or signaling.

**Figure 4.**
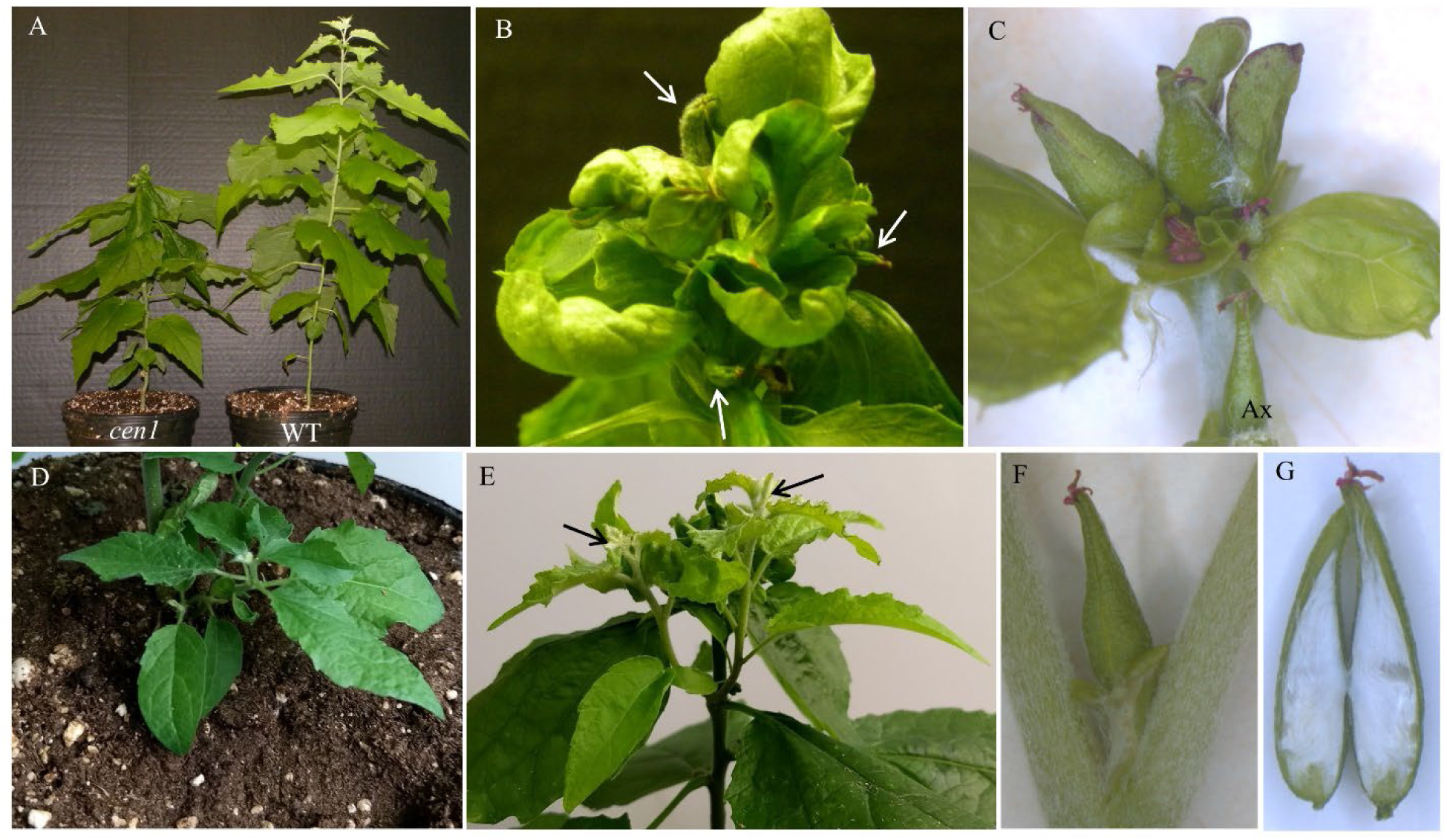
Phenotypes of potted *cen1* mutants in LDs. (A) An axillary bud of a *cen1* plant elongated to become the leader shoot and later terminated in a floral structure, whereas the WT plant main stem is continuing to grow. (B) Terminal flower structure of the *cen1* plant shown in (A). Arrows point to tops of carpels. (C) Terminal flower cluster on another *cen1* plant. Ax, axillary flower directly below the terminal flower cluster. (D) Shoot arising below the soil surface of a *cen1* plant. (E) Axillary shoots developing on a *cen1* mutant that has formed a terminal flower structure on the main stem. Arrows point to the tips of the two axillary shoots. (F) Axillary flower on a *cen1* plant and (G) bisected carpel to show cottony trichomes.

### CEN1 is necessary to prevent precocious flowering

All plants regenerated following transformation of the *CEN1*-targeting construct flowered *in vitro* (Figure S4). Six events were initially screened by direct sequencing of the PCR-amplified locus and this indicated that all had frame shift mutations. The PCR fragments from two events and WT were cloned and sequences from 10 single colonies analyzed to confirm the biallelic mutations (Figure 1, Figure S2).

All plants regenerated with the *CEN1CEN2*-targeting construct showed flowering phenotypes in tissue culture similar to the *cen1* mutants, whereas all plants with the *CEN2*-specific gRNA showed only vegetative growth (Figure S4) and we confirmed biallelic frame shift mutations in two *cen2* and two *cen1cen2* events (Figure 1, Figure S2). Shoots of most *cen1* and *cen1cen2* plants rooted and after some vegetative growth, the shoot apex terminated in a cluster of carpels and small, rounded leaves that in some cases had partial floral characteristics (Figure S4). In the axils of lower leaves, single flowers or an axillary shoot formed that recapitulated the main shoot phenotype. Carpels were often partially formed and not fully fused. This phenotype was also recapitulated by shoots that developed from roots of plants after about 3 months in tissue culture.

Two ramets each from six *cen1* mutant events were transferred to soil. All the plants had apical flower structures at the time of transplanting to soil and axillary flowers were present at some nodes. Seven plants grew to ∼15 cm in height and ceased growth, whereas the other five plants generated an axillary branch which became the leader and grew to heights ranging from 30 cm to 45 cm in height (Figure 4A). These newly formed axillary shoots initially showed only vegetative growth, but later formed apical flower structures (Figures 4B-C, S5). All these plants also formed axillary flowers (Figures 4F, S6), and in one mutant plant an axillary catkin-like structure was observed (Figure S6D). All the plants showed new shoots that originated beneath the soil surface that could have formed from the base of the stem or the roots (Figure 4D). Four *cen1* trees initiated axillary branches (Figure 4E) and both the branches and shoots sprouting from soil recapitulated the initial leader shoot phenotype. Specifically, they initially showed only vegetative growth, but then later formed terminal and axillary flowers. Many of the pistillate flowers were completely formed and produced abundant cottony seed trichomes (Figures 4G, S5C).

The *bft* mutants did not flower or show a visible phenotype compared to WT plants in tissue culture (Figure S4). *BFT* is upregulated in roots by nitrogen (N), particularly ammonium and urea, according to the Phytozome *P. trichocarpa* GeneAtlas v2 (Goodstein et al. 2012; Tuskan et al. 2006) and a previous study reported its downregulation in roots under low N (provided as KNO_3_ ;Wei et al. 2013). As an initial screen for a growth response to nutrient availability, we transferred ramets of WT and two mutant events, *bft-14* and *bft-16*, with biallelic single bp insertions (Figure S2) to two-gallon pots but did not add fertilizer. Allowing nutrients present in the potting mix to deplete leads to growth cessation and terminal bud set. We began monitoring height growth and bud stage after 104 days in soil, when all plants still had an active shoot apex (Figure S6). While the timing of growth cessation and bud set was similar in all genotypes, stem height was reduced in the *bft* mutants compared to WT (Figure S6).

### cen1 mutation induces precocious flowering in an ft1 mutant background

*FT1* is predominately expressed in winter buds, and is a potent inducer of flowering in *Populus* (Böhlenius et al. 2006; Hsu et al. 2011). In contrast, *FT*2 is expressed during the growing season and is also developmentally regulated, being strongly upregulated in leaves when they near full expansion (Hsu et al. 2011; Sheng et al. 2022). While less effective than *FT1* at inducing flowering, 35S-driven expression of *FT2* induces the formation of flowers (Hsu et al. 2011; Hsu et al. 2006). Thus, the *cen1* mutation could potentially allow either low expression of *FT1* during LD^16hr^ or *FT2* expression to induce flowering. That several leaves developed and some reached full expansion before the formation of flowers on shoots initiated on potted *cen1* plants (Figure 4), correlates with the upregulation of *FT2*, but *in vitro* plants flowered with fewer leaves. Unlike potted plants that rely on source leaves for carbon, tissue culture medium contains sugar. Thus, we compared expression of the *FTs* in a leaf gradient in tissue culture and potted plants. While *FT1* expression was relatively low in all cases, *FT2* expression showed a different developmental expression pattern in tissue culture, being highest in younger leaves (Figure 5). Finally, we introduced the construct targeting *CEN1* into the homozygous *ft1-3* mutant which exhibited WT-like growth in LD^16hr^ (Figure 2). All regenerated plants flowered *in vitro* (Figure 5) similar to the *cen1* mutants (Figure S4).

**Figure 5.**
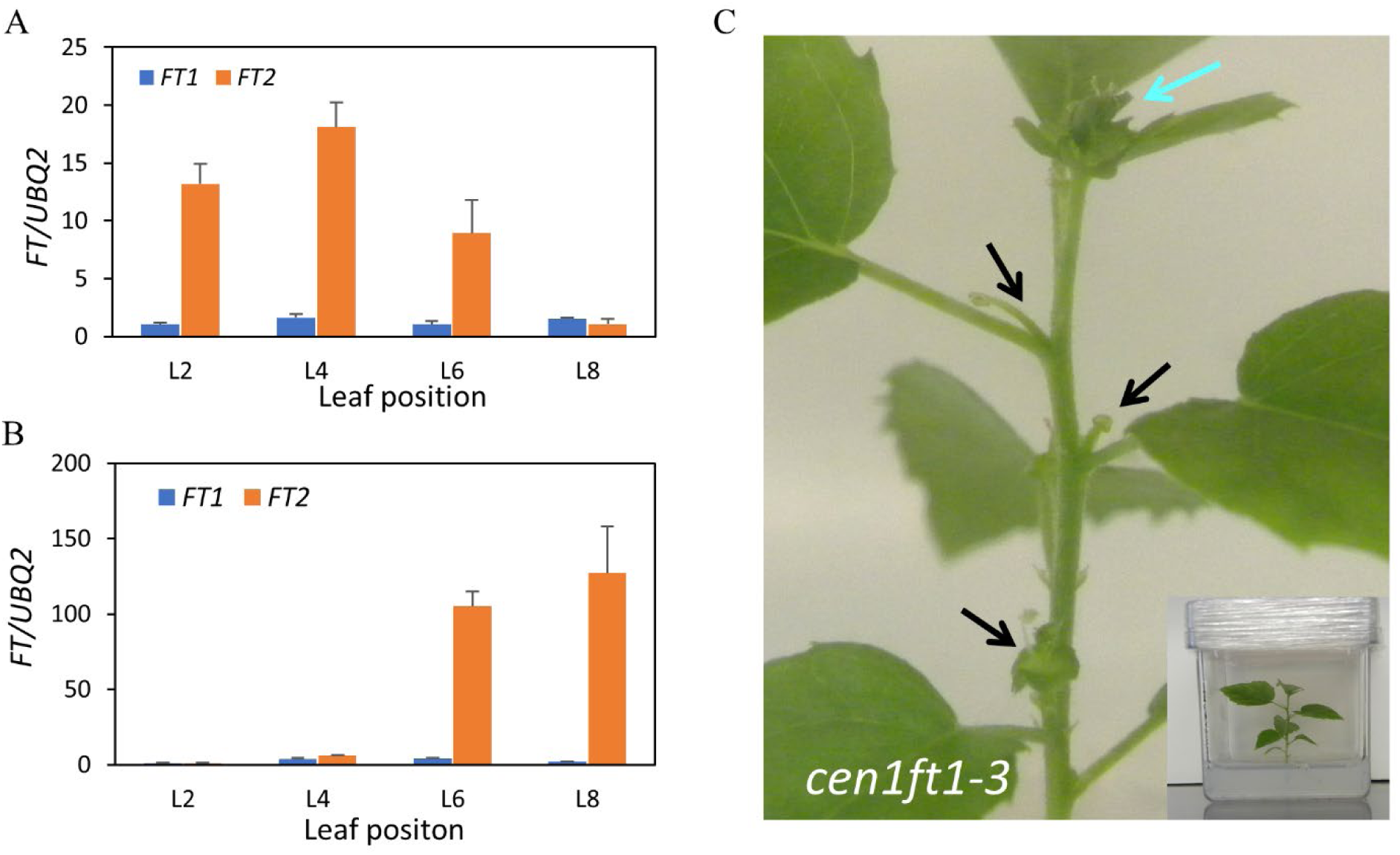
*cen1* mutation induces *in vitro* flowering in the *ft1-3* background. (A) *FT1* and *FT2* expression in a leaf gradient from an *in vitro* WT plant. (B) *FT1* and *FT2* expression in a leaf gradient from a potted WT plant. Leaf position is defined based on leaf plastochron index where leaf position 1 is the youngest leaf whose lamina is longer than 1 cm. Relative expression is fold change in *FT* transcript levels relative to the sample with the lowest expression. *FT* expression was normalized against reference gene *UBQ2*. (C) A representative *cen1ft1-3 in vitro* plant with a terminal flower structure (blue arrow) and axillary flowers (black arrows). Inset photograph shows the entire plant.

### CEN1 shows distinct seasonal expression in vegetative and reproductive tissues

The quantitative antagonism of *FT* and *CEN/TFL1* in flowering time is broadly conserved and this antagonism has been shown to extend to other processes such as potato tuber development (reviewed in Perilleux et al. 2019; Zhu et al. 2021). To be able to directly compare circannual expression of the *CEN* paralogs with the *FT* paralogs, we studied *CEN1* and *CEN2* expression in the same adult *P. deltoides* sample set previously used for study of *FT1* and *FT2* expression (Hsu et al. 2011). Whereas *CEN2* showed generally low expression in all samples, *CEN1* showed distinct seasonal patterns in all five sample types (Figure 6A, Table S3). Similar to a previous study in *P. trichocarpa* (Mohamed et al. 2010), *CEN1* expression was highly upregulated in shoot apices shortly after bud flush (Figure 6A) and though expression declined by the next month (May), expression was still relatively high until late autumn. The circannual expression profiles of *CEN1* in shoot and lateral vegetative buds (Table S3) were similar to its shoot apex profile.

**Figure 6.**
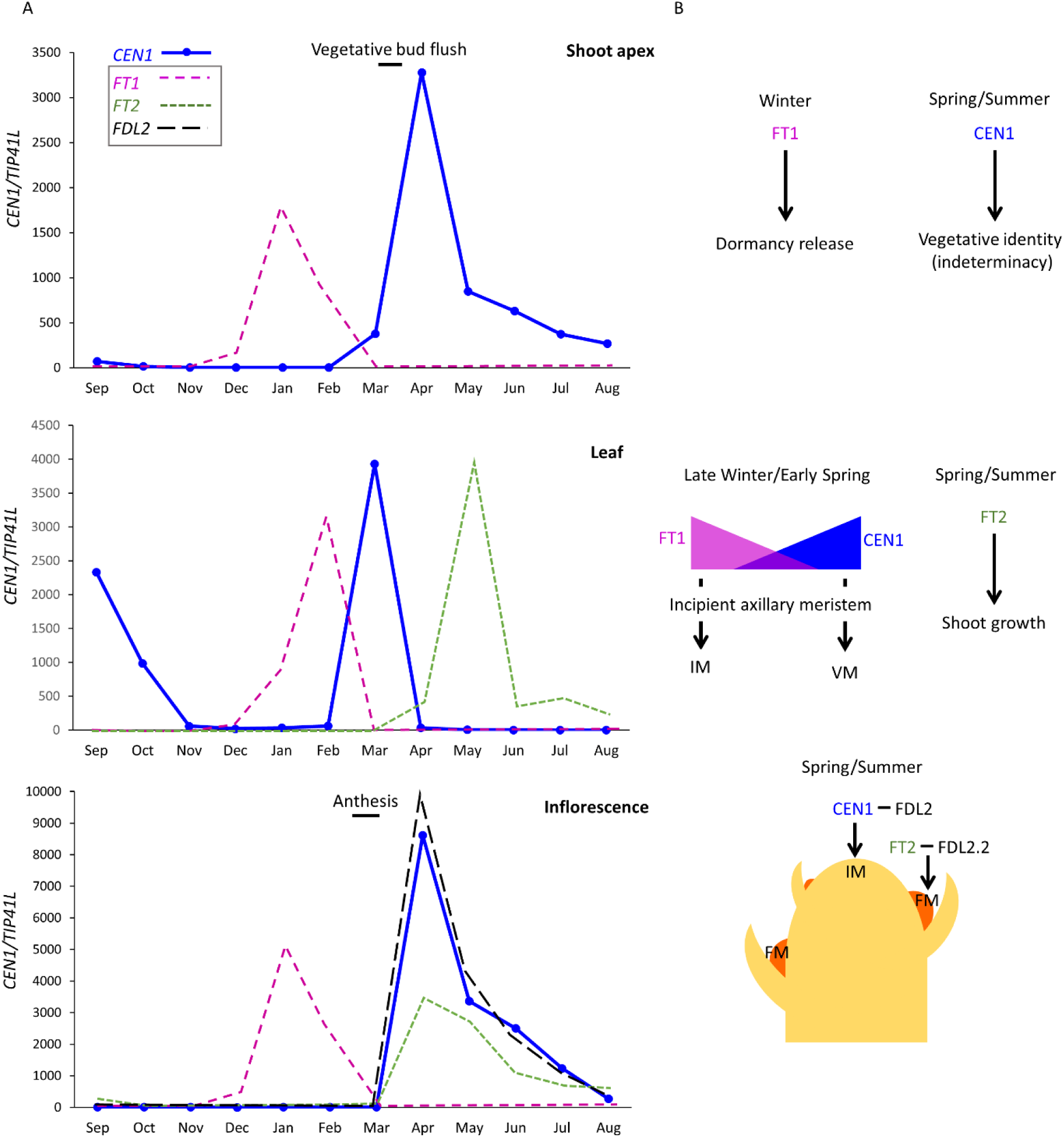
Circannual expression of *CEN1* in vegetative and reproductive tissues of adult *P. deltoides*. (A) Relative expression quantity is fold change in *CEN1* transcript levels relative to the time point with the lowest expression within a tissue (n = 3 biological replicates except for shoot apex where the apices were pooled to provide sufficient sample for analysis). *CEN1* expression was normalized against reference gene *TIP41L*. For figure clarity, standard deviations are not shown but are provided in Table S3, which also includes additional expression data. Expression of *FT1, FT2* and *FDL2* are from previous studies (Hsu et al, 2010; Sheng et al. 2022) and represent only their circannual expression patterns and not the actual fold changes that were reported. From September to March, late preformed leaves were dissected from terminal buds. Axillary inflorescence bud flush began in late February with anthesis reached in March; thus, the April sample is a newly initiated inflorescence bud. (B) Working model of *FT1, FT2*, and *CEN1* functions in the different tissues at different seasonal times. IM, inflorescence meristem; VM, vegetative meristem; FM. floral meristem.

*CEN1* expression in leaf was distinct from the other tissues, showing a strong upregulation in March LPL shortly before buds flushed, but expression rapidly declined to very low levels except for an upregulation in LPL of early autumn buds (Figure 6A, Table S3). In the vegetative sample profiles, *FT1* and *FT2* expression peaks showed little overlap with *CEN1* expression peaks except for *FT1* and *CEN1* expression in LPL before bud flush (Figure 6A; Hsu et al. 2011). *CEN1* was highly upregulated in newly initiated inflorescence buds (April) and gradually declined over the subsequent months---a circannual pattern that was highly similar to its expression in vegetative shoot apices. Interestingly, the inflorescence expression profile of *CEN1* was highly similar to the expression profile of both *FDL2* splice variants, with only *FDL2*.*2* able to promote flowering (Sheng et al. 2022). Moreover, *FT2* is also induced in initiating inflorescences at the same time (Hsu et al. 2011; Figure 6A).

## Discussion

Here we have shown that a major role for *CEN1* is to maintain indeterminate meristem identity and that *FT1* promotes dormancy release/bud flush, but the *ft1* mutation did not affect growth in LDs or SD-mediated growth cessation and bud set. We did not identify a function for *CEN2* and its low expression in the circannual sample set (Table S3) as well as in the Phytozome *P. trichocarpa* GeneAtlas v2 (Goodstein et al. 2012; Tuskan et al. 2006) does not provide clues to a function. *BFT* expression in roots is altered by N availability (Goodstein et al. 2012; Tuskan et al. 2006; Wei et al. 2013) and *bft* mutants were shorter than WT in conditions where nutrients were allowed to deplete from soil (Figure S6), suggesting that further study could reveal a role in root responses to N. *CEN1* showed distinct circannual expression patterns in vegetative and reproductive tissues and correlation with the expression patterns of *FT1* and *FT2* suggest that the relative levels of *CEN1* with *FT1* and *FT2* regulate multiple phases of a vegetative and reproductive seasonal cycles. Below we discuss the roles of these genes in different tissues at different seasonal times and the corresponding models are shown Figure 6B.

### Dormancy release and meristem indeterminacy

Study of *ft1* mutants showed that *FT1* promotes bud flush (Figure 3). Time of bud flush depends on release of dormancy by accumulation of chilling temperature units (CTUs) and subsequent accumulation of warm temperature units (WTUs) necessary to resume growth (Cooke et al. 2012). These two temperature-mediated phases are not easily separated as dormancy is dynamic; the more the accumulation of CTUs, the lower the depth of dormancy and; hence, fewer WTUs are needed to resume growth. *FT1* upregulation in January shoot apices (Figure 6a) is consistent with *FT1* promoting dormancy release, though we cannot rule out a role in warm-temperature mediated growth resumption. However, a recent study of poplar *ft1* mutants also supported that *FT1* promoted dormancy release (Andre et al. 2022a) Moreover, GA_3_ could compensate for the *ft1* mutation (Figure 3C). GA_3_ promotes dormancy release in the absence of chilling and activates a group of GH17 genes linked to the opening of plasmodesmata that are also upregulated during chilling (Rinne et al. 2011). Additionally, the very low expression of *CEN1* in winter shoot apices (Figure 6A) and that previous controlled environment study of *CEN1* overexpression and RNAi transgenics showed that *CEN1* represses dormancy release (Mohamed et al. 2010), suggests that *FT1*-mediated promotion of dormancy release depends on a high FT1:CEN1 ratio (Figure 6B).

The circannual expression profiles (Figure 6a, Table S3) and terminal flower phenotype of both *cen1* and *cen1ft1* mutants (Figures 4, 5) suggest that the marked upregulation of *CEN1* in newly flushed shoot apices is necessary to establish meristem indeterminacy before the upregulation of *FT2* in expanded leaves. Moreover, the continued expression of *CEN1* in shoot apices as well as lateral vegetative buds prevents vegetative meristems from transitioning to flowering during the growing season. Although we cannot rule out that *CEN1* inhibits other flowering promoters, the broadly conserved antagonism between *TFL1/CEN* and *FT* and that 35S promoter-driven *FT2* induces flower formation in LDs (Hsu et al. 2006), suggests that *CEN1* is preventing *FT2* from transitioning vegetative meristems to flowering during the growing season. Moreover, shoots that formed after transfer of *cen1* plants to soil, produced several phytomers and expanded leaves before producing flowers (Figure 4); thus, flowering occurred at a time when *FT2* is upregulated in expanded leaves (Figure 5B).

Indeterminacy is a property of the shoot apical meristem (SAM), which produces phytomers, while the rib meristem (RM) increases shoot elongation. Both *TFL1* and *CEN1* are expressed in the RM and presumably like TFL1, the CEN1 protein moves into the SAM to specify indeterminacy (Bradley et al. 1997; Conti and Bradley 2007; Goretti et al. 2020; Ruonala et al. 2008). *FT2* acts long distance to regulate shoot growth (Andre et al. 2022a). Although the predominant effect of *cen1* mutation on SAM determinacy might obscure effects on shoot growth, it is clear that *in vitro*-regenerated, axillary and root/stem-initiated shoots can elongate until a terminal flower is produced in *cen1* mutants (Figures 4, S4). Study of *CEN1* overexpression and RNAi transgenics in SDs indicated that it modestly promotes cessation of leaf production and bud set (Miskolczi et al. 2019; Mohamed et al. 2010). This perhaps indicates that these are mostly dependent on *FT2* expression, and rapid *FT2* downregulation in SDs (Böhlenius et al. 2006) allows *CEN1* to have some effect. This could also be the case for *FT1* in dormancy release, where in field conditions, *35S:CEN1* trees but not RNAi transgenics showed altered bud flush time compared to WT trees (Mohamed et al. 2010). Moreover, studies have not differentiated roles of the RM and SAM, and how long distance and local signals are communicated among these two meristems likely plays a key role in vegetative growth and flowering processes mediated by *FT1, FT2* and *CEN1* (Paul et al. 2014).

The roles of GA metabolism and signaling in shoot elongation and flowering are also likely central to understanding how *CEN1*-mediated SAM indeterminacy enables sustained *FT2*-mediated promotion of vegetative shoot growth but prevents *FT2*-mediated promotion of flowering during vegetative growth. GA acts both locally and systematically; grafting experiments and study of *ft2* mutants indicate that *FT2* regulates GA metabolism at the apex and mature leaf and both *FT2* and *GA20 oxidase* expression are reduced in leaf by SHORT VEGETATIVE PHASE-LIKE (Andre et al. 2022b; Gomez-Soto et al. 2021; Miskolczi et al. 2019). In Arabidopsis, increased shoot elongation (bolting) and the floral transition are coincident, and GA promotes both (reviewed in Conti 2017). However, in diverse woody angiosperms, GA promotes shoot elongation but not flowering and in some cases application of GA inhibitors is used to stimulate flowering (reviewed in Brunner et al. 2017). In *Populus*, 35S-directed expression of *GIBBERELLIC ACID INSENSITIVE*, a DELLA domain repressor of GA signaling, resulted in semi-dwarf trees with an earlier first onset of flowering, albeit catkins were upright rather than pendulous and occurred in August rather than the typical early spring (Zawaski et al. 2011).

### The transition to flowering and inflorescence development

In Arabidopsis, FT forms a transient complex with FD in the SAM to mediate its transition to an inflorescence meristem (IM) and then a TFL1-FD complex prevents the IM from transitioning to a determinate floral meristem (FM), while accumulation of FT-FD complex in new axillary meristems specifies their FM identity (reviewed in Freytes et al. 2021). In adult *Populus*, certain axillary meristems transition to inflorescence meristems that produce axillary flowers subtended by bracts. A dormant winter bud contains early preformed leaves (EPL) and LPL and detailed study of shoot development in adult *P. deltoides* showed that axillary meristems are present in EPL and that these later develop into vegetative buds (Yuceer et al. 2003); and reviewed in Brunner et al. (2014). In contrast, no meristematic domes are evident in LPL within the dormant bud, but within a few weeks following bud flush, developing inflorescence buds are distinguishable in LPL axils. Subsequent neoformed leaves (i.e., formed after bud flush) develop only vegetative buds in their axils. With the *CEN1* circannual expression profiles that can be directly compared to *FT1* profiles (Figure 6A), we propose that the shift from a high to low *FT1:CEN1* expression ratio in LPL after dormancy release but before canonical bud flush could define the seasonal window where some incipient LPL axillary meristems transition to inflorescence meristems in adult trees (Figure 6B). *TFL1* has been shown to confer an age-dependent response to vernalization in perennial *Arabis alpina*, and the effect of vernalization on *TFL1* expression differed with age (Wang et al. 2011).

Thus, it will be interesting to determine if reproductive phase change also involves changes in the expression or activity *FT1* or *CEN1* in post-dormancy LPL.

*CEN1* is highly induced in developing axillary inflorescence buds at the same time as it is upregulated in shoot apices (Figure 6A). *FT2* also shows upregulation at the same time in inflorescences as does *FDL2* (Hsu et al. 2011; Sheng et al. 2022). Mature catkins on field grown *CEN1/2*-RNAi trees had fewer flowers and were somewhat shorter than WT catkins (Mohamed et al. 2010). Considered together, these results indicate that *CEN1* also maintains indeterminacy of the IM and suggests conserved functions for *CEN1, FT2* and *FDL2* in inflorescence development (Figure 6B). Among the three *Populus FDL* genes and two *FDL2* splice variants, only *FDL2*.*2* can induce precocious flowering (Parmentier-Line and Coleman 2016; Sheng et al. 2022; Tylewicz et al. 2015). Although *FDL2*.*2* is expressed in vegetative tissues, its expression is three to four orders of magnitude greater in developing inflorescences (Sheng et al. 2022). As *CEN1* and *FT2* are also expressed in growing vegetative shoots at similar or higher levels compared to inflorescences (Figure 6A, Table S3; Hsu et al. 2011), this suggests that induction of *FDL2*.*2* expression could be a determining factor in the development of inflorescence versus vegetative shoot.

In Arabidopsis, FT promotes and TFL1 represses FM identity genes via competition for chromatin bound FD (Zhu et al. 2020). The *FDL2*.*1* splice variant encodes a protein containing an additional 29 amino acids within the conserved bZIP domain and shows circannual expression patterns similar to *FDL2*.*2* expression profiles (Sheng et al. 2022). Overexpression of WT *FDL2*.*1* or a dominant repressor version of *FDL2*.*1* induced similar dwarf phenotypes and FDL2.1 can interact with FT1 and FT2, but when co-expressed with *FT1* in rice protoplasts, it could not activate the rice *APETALA1* homolog (Sheng et al. 2022; Tylewicz et al. 2015). These results suggest that FDL2.1 could provide an additional mechanism for repressing flowering by acting as a dominant negative inhibitor via competition with FDL2.2 for interaction with FT1/2. Although considerable evidence supports that *FT1, FT2* and *FDL2*.*2* promote the floral transition and FM identity, definitive evidence of their endogenous roles is challenging due to the multi-year non-flowering phase and the key roles of *FT1* and *FT2* in vegetative growth. Nonetheless, the rapidly advancing CRISPR toolkit offers options to circumvent these challenges; for example, activation or repression rather than gene knock out and base editing to alter specific *cis* or coding sequence motifs that impact only some functions. Moreover, such *FT* mutants could be combined with CRISPR-mediated reductions in *CEN1* levels or activity that cause a more modest acceleration of flowering and allows dormancy induction/release treatments to reveal environmental interactions.

## Supplementary Data

Figure S1. Overview of CRISPR/Cas9 constructs.

Figure S2. Alignments of mutant sequences with gRNAs.

Figure S3. *ft1* axillary buds at nodes 11-23 flush the same time as WT in a detached bud-internode assay.

Figure S4. Phenotypes of *cen1, cen2, cen1cen2* and *bft* plants in tissue culture.

Figure S5. Axillary flowers on potted *cen1* trees that had produced terminal flower structures.

Figure S6. *bft* mutants show reduced height growth compared to WT, but they cease growth and set terminal buds similar to WT as nutrients deplete.

Table S1. Primer sequences and gene IDs.

Table S2. Expression data for *FT1* and *FT2* in *P. tremula x P. alba* INRA 717-1B.

Table S3. Seasonal expression data for *CEN1* and *CEN2* in adult *P. deltoides*.

## Authors’ Contributions

XS, RAM and C-TW designed and executed the experimental work and analyzed data. AMB directed the research, analysed results and drafted the paper with XS.

## References

Andre, D., A. Marcon, K.C. Lee, D. Goretti, B. Zhang, N. Delhomme, M. Schmid and O. Nilsson. 2022a. FLOWERING LOCUS T paralogs control the annual growth cycle in Populus trees. Current Biology. 32:2988-+.

Andre, D., J.A. Zambrano, B. Zhang, K.C. Lee, M. Ruhl, A. Marcon and O. Nilsson. 2022b. Populus SVL Acts in Leaves to Modulate the Timing of Growth Cessation and Bud Set. Front Plant Sci. 13:823019.

Bennett, T. and L.E. Dixon. 2021. Asymmetric expansions of FT and TFL1 lineages characterize differential evolution of the EuPEBP family in the major angiosperm lineages. BMC Biol. 19:181.

Böhlenius, H., T. Huang, L. Charbonnel-Campaa, A.M. Brunner, S. Jansson, S.H. Strauss and O. Nilsson. 2006. CO/FT regulatory module controls timing of flowering and seasonal growth cessation in trees. Science. 312:1040–3.

Bradley, D., O. Ratcliffe, C. Vincent, R. Carpenter and E. Coen. 1997. Inflorescence commitment and architecture in Arabidopsis. Science. 275:80–3.

Brunner, A.M., L.M. Evans, C.-Y. Hsu and X. Sheng. 2014. Vernalization and the Chilling Requirement to Exit Bud Dormancy: Shared or Separate Regulation? Frontiers in Plant Science. 5

Brunner, A.M., E. Varkonyi-Gasic and R.C. Jones. 2017. Phase Change and Phenology in Trees. In Comparative and Evolutionary Genomics of Angiosperm Trees Eds. A. Groover and Q. Cronk. Springer, Cham, Switzerland, pp 227–274.

Brunner, A.M., I.A. Yakovlev and S.H. Strauss. 2004. Validating internal controls for quantitative plant gene expression studies. BMC Plant Biology. 4:14.

Conti, L. 2017. Hormonal control of the floral transition: Can one catch them all? Developmental Biology. 430:288–301.

Conti, L. and D. Bradley. 2007. TERMINAL FLOWER1 is a mobile signal controlling Arabidopsis architecture. Plant Cell. 19:767–778.

Cooke, J.E., M.E. Eriksson and O. Junttila. 2012. The dynamic nature of bud dormancy in trees: environmental control and molecular mechanisms. Plant Cell Environ. 35:1707–28.

Eshed, Y. and Z.B. Lippman. 2019. Revolutions in agriculture chart a course for targeted breeding of old and new crops. Science. 366

Evans, L.M., G.T. Slavov, E. Rodgers-Melnick, J. Martin, P. Ranjan, W. Muchero, A.M. Brunner, W. Schackwitz, L. Gunter, J.G. Chen, G.A. Tuskan and S.P. DiFazio. 2014. Population genomics of Populus trichocarpa identifies signatures of selection and adaptive trait associations. Nature Genetics. 46:1089–1096.

Freytes, S.N., M. Canelo and P.D. Cerdan. 2021. Regulation of Flowering Time: When and Where? Current Opinion in Plant Biology. 63

Gomez-Soto, D., I. Allona and M. Perales. 2021. FLOWERING LOCUS T2 Promotes Shoot Apex Development and Restricts Internode Elongation via the 13-Hydroxylation Gibberellin Biosynthesis Pathway in Poplar. Front Plant Sci. 12:814195.

Goodstein, D.M., S. Shu, R. Howson, R. Neupane, R.D. Hayes, J. Fazo, T. Mitros, W. Dirks, U. Hellsten, N. Putnam and D.S. Rokhsar. 2012. Phytozome: a comparative platform for green plant genomics. Nucleic Acids Research. 40:D1178–D1186.

Goretti, D., M. Silvestre, S. Collani, T. Langenecker, C. Méndez, F. Madueño and M. Schmid. 2020. TERMINAL FLOWER1 Functions as a Mobile Transcriptional Cofactor in the Shoot Apical Meristem1 [OPEN]. Plant Physiology. 182:2081–2095.

Gutierrez, L., M. Mauriat, S. Guenin, J. Pelloux, J.F. Lefebvre, R. Louvet, C. Rusterucci, T. Moritz, F. Guerineau, C. Bellini and O. Van Wuytswinkel. 2008. The lack of a systematic validation of reference genes: a serious pitfall undervalued in reverse transcription-polymerase chain reaction (RT-PCR) analysis in plants. Plant Biotechnol J. 6:609–18.

Hsu, C.Y., J.P. Adams, H. Kim, K. No, C. Ma, S.H. Strauss, J. Drnevich, L. Vandervelde, J.D. Ellis, B.M. Rice, N. Wickett, L.E. Gunter, G.A. Tuskan, A.M. Brunner, G.P. Page, A. Barakat, J.E. Carlson, C.W. DePamphilis, D.S. Luthe and C. Yuceer. 2011. FLOWERING LOCUS T duplication coordinates reproductive and vegetative growth in perennial poplar. Proc Natl Acad Sci U S A. 108:10756–61.

Hsu, C.Y., Y. Liu, D.S. Luthe and C. Yuceer. 2006. Poplar FT2 shortens the juvenile phase and promotes seasonal flowering. Plant Cell. 18:1846–61.

Jin, S., Z. Nasim, H. Susila and J.H. Ahn. 2021. Evolution and functional diversification of FLOWERING LOCUS T/TERMINAL FLOWER 1 family genes in plants. Semin Cell Dev Biol. 109:20–30.

Klocko, A.L., E. Borejsza-Wysocka, A.M. Brunner, O. Shevchenko, H. Aldwinckle and S.H. Strauss. 2016. Transgenic Suppression of AGAMOUS Genes in Apple Reduces Fertility and Increases Floral Attractiveness. Plos One. 11

Larson, P.R. and J.G. Isebrands. 1971. The Plastochron Index as Applied to Developmental Studies of Cottonwood. Canadian Journal of Forest Research. 1:1–11.

Lawrence, E.H., A.R. Leichty, E.E. Doody, C. Ma, S.H. Strauss and R.S. Poethig. 2021. Vegetative phase change in Populus tremula × alba. New Phytol. 231:351–364.

Li, J.F., J.E. Norville, J. Aach, M. McCormack, D.D. Zhang, J. Bush, G.M. Church and J. Sheen. 2013. Multiplex and homologous recombination-mediated genome editing in Arabidopsis and Nicotiana benthamiana using guide RNA and Cas9. Nature Biotechnology. 31:688–691.

Livak, K.J. and T.D. Schmittgen. 2001. Analysis of Relative Gene Expression Data Using Real-Time Quantitative PCR and the 2−ΔΔCT Method. Methods. 25:402–408.

Meilan, R. and C. Ma. 2006. Poplar (Populus spp.). Methods in molecular biology (Clifton, N.J.). 344:143–151.

Miskolczi, P., R.K. Singh, S. Tylewicz, A. Azeez, J.P. Maurya, D. Tarkowska, O. Novak, K. Jonsson and R.P. Bhalerao. 2019. Long-range mobile signals mediate seasonal control of shoot growth. Proc Natl Acad Sci U S A. 116:10852–10857.

Mohamed, R., C.T. Wang, C. Ma, O. Shevchenko, S.J. Dye, J.R. Puzey, E. Etherington, X. Sheng, R. Meilan, S.H. Strauss and A.M. Brunner. 2010. Populus CEN/TFL1 regulates first onset of flowering, axillary meristem identity and dormancy release in Populus. Plant J. 62:674–88.

Parmentier-Line, C.M. and G.D. Coleman. 2016. Constitutive expression of the Poplar FD-like basic leucine zipper transcription factor alters growth and bud development. Plant Biotechnology Journal. 14:260–270.

Paul, L.K., P.L.H. Rinne and C. van der Schoot. 2014. Shoot meristems of deciduous woody perennials: self-organization and morphogenetic transitions. Current Opinion in Plant Biology. 17:86–95.

Perilleux, C., F. Bouche, M. Randoux and B. Orman-Ligeza. 2019. Turning Meristems into Fortresses. Trends in Plant Science. 24:431–442.

Peterson, B.A., D.C. Haak, M.T. Nishimura, P.J. Teixeira, S.R. James, J.L. Dangl and Z.L. Nimchuk. 2016. Genome-Wide Assessment of Efficiency and Specificity in CRISPR/Cas9 Mediated Multiple Site Targeting in Arabidopsis. PLoS One. 11:e0162169.

Rinne, P.L., A. Welling, J. Vahala, L. Ripel, R. Ruonala, J. Kangasjarvi and C. van der Schoot. 2011. Chilling of dormant buds hyperinduces FLOWERING LOCUS T and recruits GA-inducible 1,3-beta-glucanases to reopen signal conduits and release dormancy in Populus. Plant Cell. 23:130–46.

Ruonala, R., P.L.H. Rinne, J. Kangasjarvi and C. van der Schoot. 2008. CENL1 expression in the rib meristem affects stem elongation and the transition to dormancy in Populus. Plant Cell. 20:59–74.

Saure, M.C. 1985. Dormancy Release in Deciduous Fruit Trees. In Horticultural Reviews.

Sheng, X., C.Y. Hsu, C. Ma and A.M. Brunner. 2022. Functional Diversification of Populus FLOWERING LOCUS D-LIKE3 Transcription Factor and Two Paralogs in Shoot Ontogeny, Flowering, and Vegetative Phenology. Front Plant Sci. 13:805101.

Tuskan, G.A., S. Difazio, S. Jansson, J. Bohlmann, I. Grigoriev, U. Hellsten, N. Putnam, S. Ralph, S. Rombauts, A. Salamov, J. Schein, L. Sterck, A. Aerts, R.R. Bhalerao, R.P. Bhalerao, D. Blaudez, W. Boerjan, A. Brun, A. Brunner, V. Busov, M. Campbell, J. Carlson, M. Chalot, J. Chapman, G.L. Chen, D. Cooper, P.M. Coutinho, J. Couturier, S. Covert, Q. Cronk, R. Cunningham, J. Davis, S. Degroeve, A. Dejardin, C. Depamphilis, J. Detter, B. Dirks, I. Dubchak, S. Duplessis, J. Ehlting, B. Ellis, K. Gendler, D. Goodstein, M. Gribskov, J. Grimwood, A. Groover, L. Gunter, B. Hamberger, B. Heinze, Y. Helariutta, B. Henrissat, D. Holligan, R. Holt, W. Huang, N. Islam-Faridi, S. Jones, M. Jones-Rhoades, R. Jorgensen, C. Joshi, J. Kangasjarvi, J. Karlsson, C. Kelleher, R. Kirkpatrick, M. Kirst, A. Kohler, U. Kalluri, F. Larimer, J. Leebens-Mack, J.C. Leple, P. Locascio, Y. Lou, S. Lucas, F. Martin, B. Montanini, C. Napoli, D.R. Nelson, C. Nelson, K. Nieminen, O. Nilsson, V. Pereda, G. Peter, R. Philippe, G. Pilate, A. Poliakov, J. Razumovskaya, P. Richardson, C. Rinaldi, K. Ritland, P. Rouze, D. Ryaboy, J. Schmutz, J. Schrader, B. Segerman, H. Shin, A. Siddiqui, F. Sterky, A. Terry, C.J. Tsai, E. Uberbacher, P. Unneberg, et al. 2006. The genome of black cottonwood, Populus trichocarpa (Torr. & Gray). Science. 313:1596–604.

Tylewicz, S., H. Tsuji, P. Miskolczi, A. Petterle, A. Azeez, K. Jonsson, K. Shimamoto and R.P. Bhalerao. 2015. Dual role of tree florigen activation complex component FD in photoperiodic growth control and adaptive response pathways. Proc Natl Acad Sci U S A. 112:3140–5.

Wang, J., J.H. Ding, B.Y. Tan, K.M. Robinson, I.H. Michelson, A. Johansson, B. Nystedt, D.G. Scofield, O. Nilsson, S. Jansson, N.R. Street and P.K. Ingvarsson. 2018. A major locus controls local adaptation and adaptive life history variation in a perennial plant. Genome Biology. 19

Wang, R., M.C. Albani, C. Vincent, S. Bergonzi, M. Luan, Y. Bai, C. Kiefer, R. Castillo and G. Coupland. 2011. Aa TFL1 Confers an Age-Dependent Response to Vernalization in Perennial Arabis alpina The Plant Cell. 23:1307–1321.

Wei, H.R., Y.S. Yordanov, T. Georgieva, X. Li and V. Busov. 2013. Nitrogen deprivation promotes Populus root growth through global transcriptome reprogramming and activation of hierarchical genetic networks. New Phytologist. 200:483–497.

Wickland, D.P. and Y. Hanzawa. 2015. The FLOWERING LOCUS T/TERMINAL FLOWER 1 Gene Family: Functional Evolution and Molecular Mechanisms. Mol Plant. 8:983–97.

Yuceer, C., S.B. Land, M.E. Kubiske and R.L. Harkess. 2003. Shoot morphogenesis associated with flowering in Populus deltoides (Salicaceae). American Journal of Botany. 90:196–206.

Zawaski, C., M. Kadmiel, J. Pickens, C. Ma, S. Strauss and V. Busov. 2011. Repression of gibberellin biosynthesis or signaling produces striking alterations in poplar growth, morphology, and flowering. Planta. 234:1285–1298.

Zhu, Y., S. Klasfeld, C.W. Jeong, R. Jin, K. Goto, N. Yamaguchi and D. Wagner. 2020. TERMINAL FLOWER 1-FD complex target genes and competition with FLOWERING LOCUS T. Nat Commun. 11:5118.

Zhu, Y., S. Klasfeld and D. Wagner. 2021. Molecular regulation of plant developmental transitions and plant architecture via PEPB family proteins: an update on mechanism of action. Journal of Experimental Botany. 72:2301–2311.

