## Supplemental Figures 1-6 for "CRISPR/Cas9 mutants delineate roles of *Populus FT* and *TFL1*/*CEN*/*BFT* family members in growth, dormancy release and flowering"

**Supplementary Figures**

A

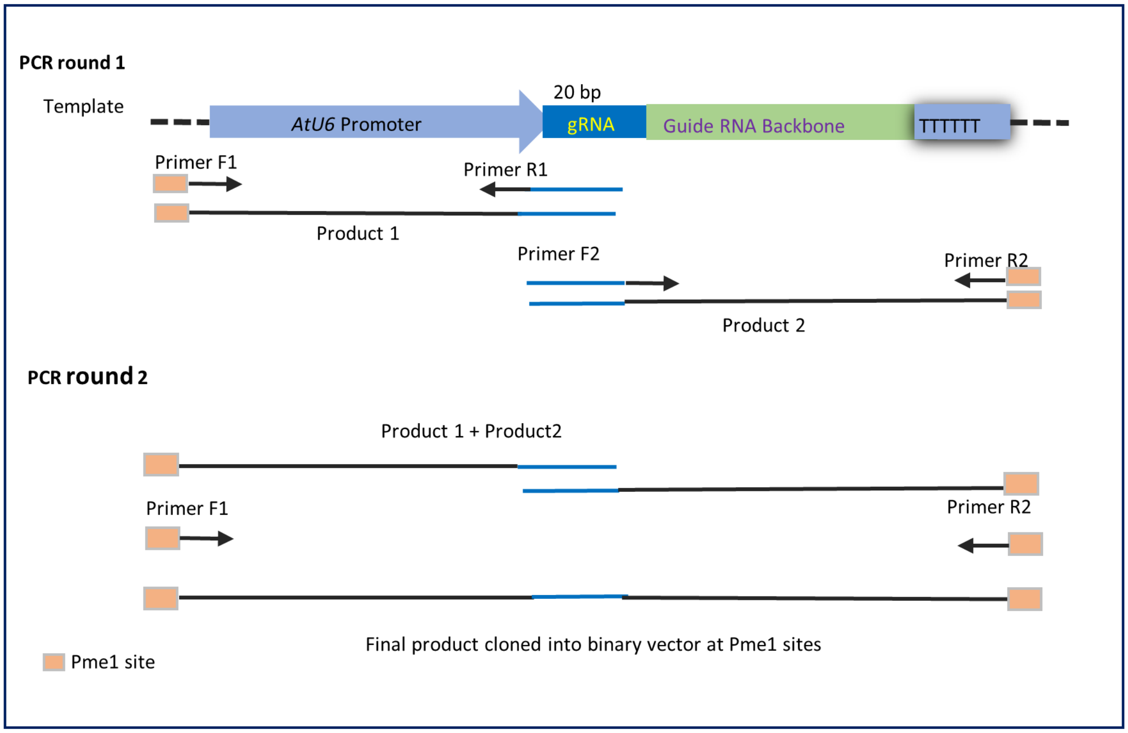


B

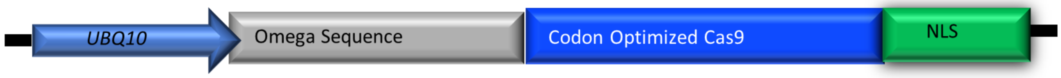


Figure S1. Overview of CRISPR/Cas9 constructs. A. Construction of gRNA cassettes. B. Features of the Cas9 transgene inserted into the pMOA33 binary vector (Peterson et al. 2016). The TMV omega translational enhancer was inserted between the Arabidopsis UBQ10 promoter and the dual maize and Arabidopsis optimized Cas9 coding sequence, which ends with an OCS terminator and is fused to the N7 nuclear localization sequence (NLS).

| Genotype | Gene/Allele | Target site sequence |
|---|---|---|
| | | GAGAGCCCGAGGCCGA-CGATGG |
| *ft1-1* | *FT1allele1* | TATGAGAGCCCGAGGCCGA---ATGGGGA |
| | *FT1allele2* | TATGAGAGCCCGAGGCCGATCGATGGGGA |
| *ft1-3* | *FT1* | TATGAGAGCCCGAGGCCGAACGATGGGGA |
| | | CTAAGGACCTTCTACA-CTCTGG |
| *ft1ft2-4* | *FT1* | GATCTAAGGACCTTCTACAACTCTGGTACGTCT |
| | *FT2* | GATCTAAGGACCTTCTAC--CTCTGGTAAGTCC |
| *ft1ft2-13* | *FT1* | GATCTAAGGACCTTCTACATCTCTGGTACGTCT |
| | *FT2* | GATCTAAGGACCTTCTAC--CTCTGGTAAGTCC |
| *ft1ft2-20* | *FT1* | GATCTAAGGACCTTCTAC--CTCTGGTACGTCT |
| | *FT2* | GATCTAAGGACCTTCTAC--CTCTGGTAAGTCC |
| | | AAAGATGTCAGAGCCTC--TTGTGG |
| *cen1-1* | *CEN1allele 1* | GGCAAAGATGTCAGAGCCTCTTTTGTGGTTG |
|  | *CEN1allele 2* | GGCAAAGATGTCAGAGCC----TTGTGGTTG |
| *cen1-4* | *CEN1* | GGCAAAGATGTCAGAGCCT---TTGTGGTTG |
| | | CCTAAAGTTGAGGTCCATGGTGG |
| *cen2-1* | *CEN2* | AAACCTAAAGTTGAGGTCC-TGGTGGTGA |
| *cen2-3* | *CEN2* | AAACCTAAAGTTGAGGTCC-TGGTGGTGA |
| | | CTTGTGGTTGGGAGAGT-GATTGG |
| *cen1cen2-1* | *CEN1allele1* | AGAGCCTCTTGTGGTTGGGAGAGTTGATTGGAGATGTT |
| | *CEN1allele2* | AGAGCCTCTTGTGGTTGGGAGAGTAGATTGGAGATGTT |
| | *CEN2allele1* | GGATCCTCTTGTGGTTGGGAGAGTTGATTGGAGATGTA |
| | *CEN2allele2* | GGATCCTCTTGTGGTTGGGAGAGT--ATTGGAGATGTA |
| *cen1cen2-4* | *CEN1* | AGAGCCTCTTGTGGTTGGGAGAGTTGATTGGAGATGTT |
| | *CEN2* | GGATCCTCTTGTGGTTGGGAGAGTTGATTGGAGATGTA |
| | | TAGAGTTGAGATTGGTG-GTGAGG |
| *bft-14* | *BFT* | ACCTAGAGTTGAGATTGGTGGGTGAGGACA |
| *bft-16* | *BFTallele1* | ACCTAGAGTTGAGATTGGTGGGTGAGGACA |
| | *BFTallele2* | ACCTAGAGTTGAGATTGGTGTGTGAGGACA |

Figure S2. Alignments of mutant sequences with gRNAs. Sequences were obtained from 10 independently cloned PCR-amplified fragments for each mutant. All mutants are biallelic and if mutants are heterozygous, sequences for both alleles are shown. Mutations are indicated by red type in yellow background and the gRNA is in blue type with PAM underlined. In double mutants, sequence differences among paralogs are in pink type. As all plants regenerated following the transformation of the *ft1-3* mutant with the construct targeting *CEN1* were phenocopies (flowered *in vitro*) of the *cen1* mutants, we did not sequence the *CEN1* target locus in the *cen1ft1-3* mutants.

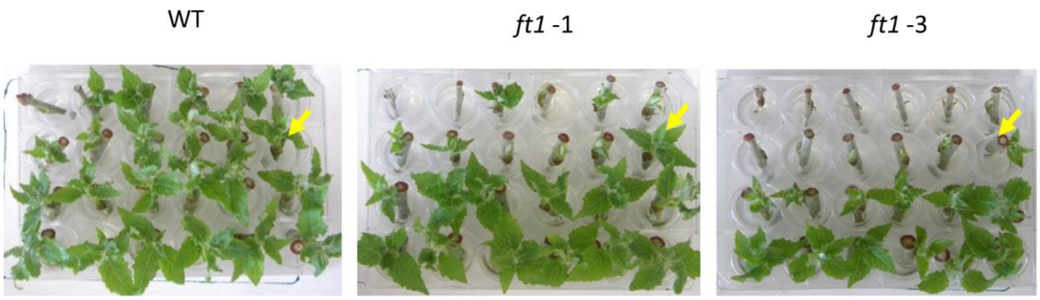


Figure S3. *ft1* axillary buds at nodes 11-23 flush the same time as WT in a detached bud-internode assay. Trees (n=4 per genotype) were exposed to 8 weeks of SDs followed by 6 weeks of SD/chilling temperatures. Stems were then cut into bud-internode units and placed in water in 24-well plates at room temperature and constant light to promote bud flush. Buds are ordered the same in each plate with the terminal bud unit in the upper left well and then successive axillary bud units ordered left to right, top to bottom. Arrows point to the axillary bud at node 11. Photos were taken two weeks after bud units were placed in water.

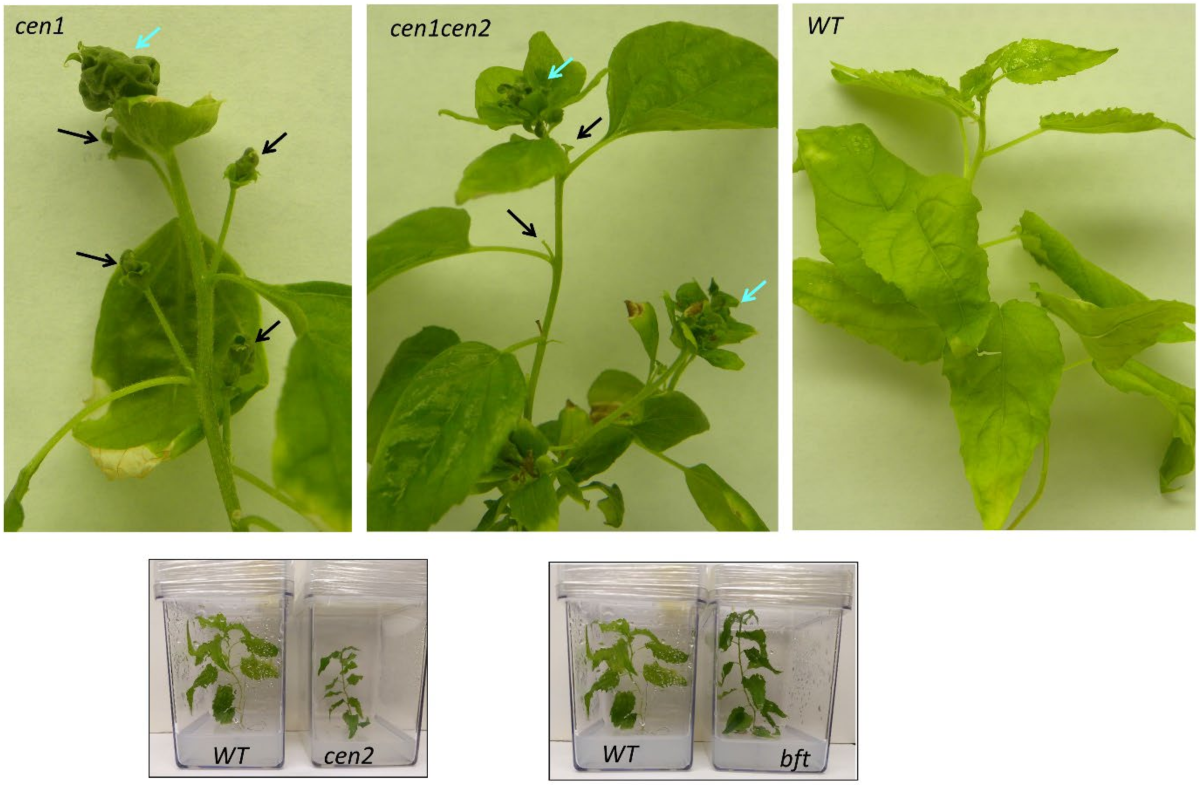


Figure S4. Phenotypes of *cen1*, *cen2* , *cen1cen2* and *bft* plants in tissue culture. Blue arrows point to terminal flower structures and black arrows point to axillary flowers on *cen1* and *cen1 cen2* mutants. In some cases, axillary flowers are incomplete or most floral parts have abscised.

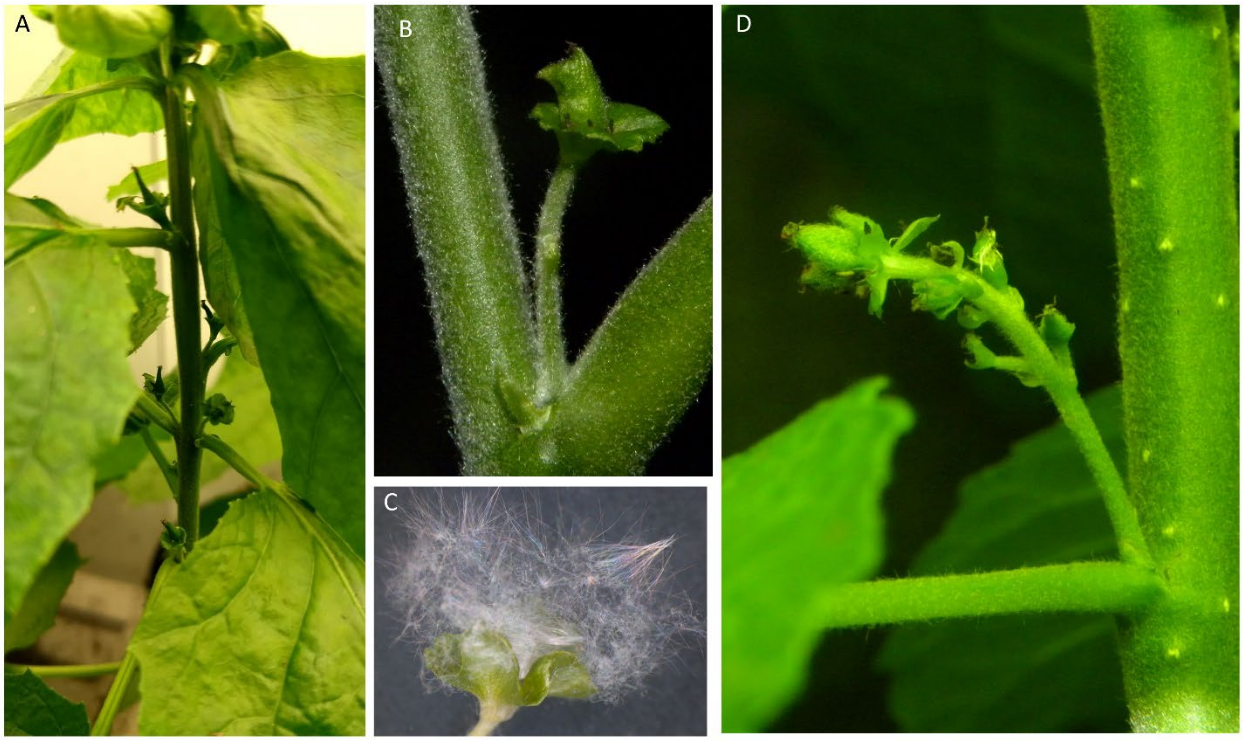


Figure S5. Axillary flowers on potted *cen1* trees that had produced terminal flower structures.

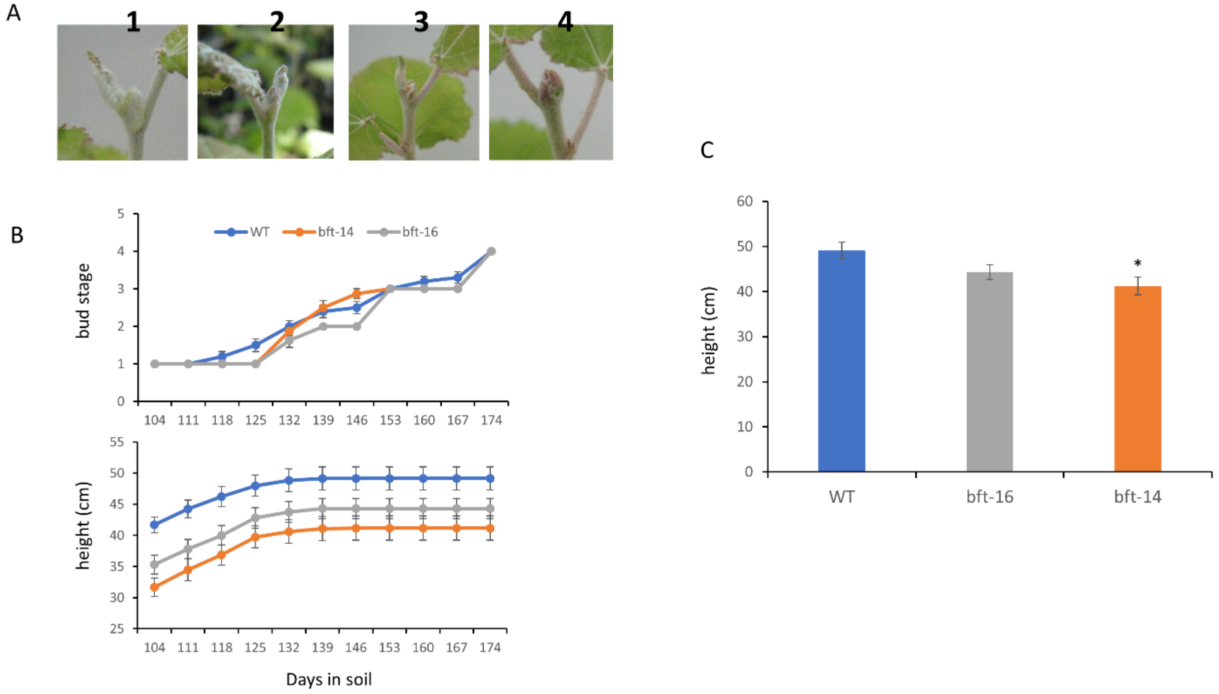


Figure S6. *bft* mutants show reduced height growth compared to WT, but they cease growth and set terminal buds similar to WT as nutrients deplete. A) Bud stage scoring system: 1- fully active apex, more than two curled up leaves; 2-bud scales are visible and extend part way up apex; 3-predominantly green bud scales cover shoot apex; 4-bud set, bud scales are predominantly reddish brown. B) Progression of bud stage and C) growth cessation in WT and *bft* mutant events. D) Final mean height of WT and *bft* mutant events. Means ± SE, n =10 for WT and n=8 per each *bft* mutant event. *significantly different from WT, student's t-test, p=0.013
